# Exploring thermal tolerance across time and space in a tropical bivalve, *Pinctada margaritifera*

**DOI:** 10.1101/2024.04.02.587172

**Authors:** K. Lugue, C. J. Monaco, E. Vigouroux, M. Sham Koua, J. Vidal-Dupiol, G. Mitta, J. Le Luyer

**Affiliations:** UMR-241 SECOPOL, Ifremer, IRD, Institut Louis-Malardé, Univ. Polynésie Française, F-98725 Taravao, Tahiti, Polynésie Française, France; IHPE, Ifremer, Univ. Montpellier, CNRS, Univ. Perpignan Via Domitia, Montpellier, France; Ifremer, Univ Brest, CNRS, IRD, UMR 6539, LEMAR, F-29280, Plouzané, France

**Keywords:** thermal tolerance, thermal limits, thermal death time, critical temperature, tropical bivalve, global warming vulnerability, ontogeny

## Abstract

Ectotherm’s vulnerability to climate change is predicted to increase with temperature variations. Still, translating laboratory observations of organisms’ heat stress responses to the natural fluctuating environment remains challenging. In this study, we used an integrative framework combining insights from the TDT curves and physiological reaction norms, to precisely capture *Pinctada margaritifera* stage -specific thermal tolerance. On a second hand, this study aimed to explore applicability of the model of accumulation of thermal injury, by making in situ predictions at three contrasting sites across French Polynesia. By working with two life stages (early planktonic vs. spat), our study revealed an ontogenetic shift in lethal (CTmax) and sub-lethal (Tc) thermal limits, with higher vulnerability for early larval stages. Cumulative injury calculations resulted in thermal failure (100% injury) for larvae within 12 and 22 h, at the most ‘extreme’ site (Nuku Hiva; T°C > 38°C), and warm lagoon (Reao Atoll), respectively. While substantial damages accumulated in spats, when exposed to consecutive extreme tides (Nuku Hiva) for 8 days. Overall, our results revealed that *P. margarifiera* is living closer to their upper thermal limits than previously estimated, and inhabit environments where important reduction of settlement and heat stress are already occurring during warmest months.

## 1. Introduction

Temperature is one of the most pervasive drivers of species eco-evolutionary dynamics. Not surprisingly, an extensive body of literature exists trying to answer the questions: (i) How do organisms perform across the range of temperatures they experience currently, and assuming climate-change scenario projections (e.g., Sinclair et al., 2016a)?, and (ii) what is the future of natural populations and ecosystems in the face of ongoing rapid climate change (e.g., Pigot et al., 2023) ?

Thermal performance curves (TPCs), which describe the performance of fitness-related traits as a function of body temperature (Gilchrist, 1995; Huey & Kingslover, 1989; Huey & Stevenson, 1979), are used extensively as an heuristic tool to predict ectothermic organism’s responses to climate change (Angilletta, 2009; Little & Seebacher, 2021; Sinclair et al., 2016b). Indeed, this unimodal and asymmetric modelling curve provides information on how specific physiological systems react to temperature as well as critical thermal limits (e.g., the temperature at which performance is zero; CTmin and CTmax). Thermal tolerance limits in particular, have become a fundamental proxy to assess species vulnerability to climate change (e.g., Sunday et al., 2012). Importantly, however, the insights gathered from both TPCs and critical thermal limits can depend strongly on the methods used (Chown et al., 2009; Kellermann et al., 2019), thus complicating its application, particularly for fluctuating environments (Khelifa et al., 2019; Kingsolver et al., 2015). Consequently, there is little consensus on the ideal method for estimating thermal limits, which limits comparisons across studies and engenders confusion and misuse of terminology (Clusella-Trullas et al., 2021; Cooper et al., 2008; Hoffmann et al., 2003).

To explicitly account for the variability in intensity and duration of thermal stress experienced by organisms in nature, Rezende et al. (2014) have recently formalised the concept of the thermal death time (TDT). Building from a rich history of earlier studies that recognized the importance of exposure duration on the estimation of ectotherms’ thermal limits (Ansell et al., 1981; Bigelow, 1921; Coles et al., s. d.; Foster, 1969; Fry et al., 1946; Kilgour & McCauley, 1986; Nedved et al., 1998; Smith, 1957; Urban, 1994), the TDT framework provides a standardized method to reconcile static and dynamic experimental assays (Jørgensen et al., 2021a, 2021a), allowing generalized comparisons across species (Molina et al., 2023; Vives□Ingla et al., 2023; Willot et al., 2022), populations (Castañeda et al., 2015; Li et al., 2023), life stage (Truebano et al., 2018), life-history traits, including prior acclimation (Tarapacki et al., 2021) or nutrient quality treatments (Koussoroplis et al., 2023).

Moreover, the integration of TDT and TPC models (Ørsted et al., 2022a) provides a consistant and global picture of organism’s thermal niche. But most importantly, this framework enables the identification of the critical temperature (Tc), i.e. at the very extrem edge of the *permissive* temperature range (i.e., possible life completion). Once environmental temperatures (Tenv) exceeds this sublethal temperature (Tc); and the *stressful* temperature range crossed (i.e., accumulation heat failure); it becomes possible to predict mortality rates under variable thermal regimes (Jørgensen et al., 2021a), and offers possibilities to more detailed, mechanistic explorations at the molecular level.

Tropical marine ectotherms are expected to be most severely affected by global warming (Deutsch et al., 2008; Dillon et al., 2010; Jørgensen et al., 2022; Nguyen et al., 2011; Pinsky et al., 2019; Tewksbury et al., 2008). Still, current assessments of thermal performance of tropical marine species are incomplete, focusing mostly on the adult life stage, and thus ignoring the thermal sensitivity of more vulnerable early-life stages (Dahlke et al., 2020). This finding is particularly worrying for tropical marine ectotherms with early dispersive stages, governing population abundances and dynamics. In French Polynesia, pearl production is essentially based on settlement of pelagic larva of *Pinctada margaritifera*, on immersed supports (i.e. spat collectors; Southgate & Lucas, 2008). While extensive work has been done for this socio-economically important species; including the calibration of a Dynamic Energy Model (DEB) for growth, survival, reproduction and larval development (Sangare et al., 2019, 2020; Thomas et al., 2011); the upper thermal limits of the early sensitive life stages remain largely uncharacterised.

To fill this knowledge gap, we used an integrative framework combining insights from the TDT curves and physiological reaction norms derived from organisms’ energy budget data, to precisely capture *P. margaritifera* stage -specific critical temperature (Tc). By providing these quantitative means, this study aimed to make *in situ* predictions of accumulation of thermal injury across time and space, and establish a necessary foundation to further our understanding of species resilience (acclimatization and genetic-adaptation potential) to climate warming trends and punctual extreme events.

## 2. Materials and methods

Three different experiments (E1, E2 and E3, details below) were performed using hatchery-produced individuals issued from two reproduction events (see cohorts A and B in **Supplementary information**). The E1 experiment aimed to assess larval development completion (i.e., performance and upper limits) under a wide range of temperatures and intensities (i.e. time of exposure). In turn, the E2 and E3 allowed investigating the thermal sensitivity of spats experiencing stressful and permissive temperatures, respectively.

### 2.1 Thermal stress assays

#### 2.1.1 Upper limits and thermal performance of larvae (E1)

This experiment had a two-fold objective: (i) describe thermal performance breath during the embryogenesis (i.e., 1-6 hours post fertilization [hpf]) and pelagic larval development phases (i.e., 1-24 hpf), and (ii) build a TDT curve for the pelagic larval development. To achieve these objectives, a full-factorial experiment was performed, including seven temperatures (22, 26, 30, 32, 34, 36, and 38 °C) x two exposure durations (6 and 24 h), as well as a control (28 °C - 24 h). In order to buid a TDT curve, two exposure durations (3 and 12 h) were additionnaly investigated for *warm* temperature conditions (i.e. > 28 °C). We chose 24 h as the longest exposure duration based on the time taken by embryos to develop into straight-hinge D-shaped larvae, with completely developed shells, as established in the laboratory (Doroudi & Southgate, 2003). Each combinaition was run in triplicate. We used one-hpf zygotes that were incubated in 15-mL Falcon^®^ tubes containing 10 mL of 10-μm filtered, UV-treated seawater at ∼28 °C (density per tube ∼ 30 zygotes mL^-1^) immersed in temperature-controlled experimental tanks. The zygotes were exposed to a fast warming/cooling (1.5 °C.min^1^), reaching their respective treatment temperatures within 10 minutes. After the treatment period, each sample was preserved by adding 100 μL of formol. The D-shaped larvae were counted and photographed using a LEICA H80 stereo microscope.

#### 2.1.2 Upper thermal limits of juveniles (E2)

We examined the effect of 11 *warm* temperatures, ranging from 32 to 42 °C, on the survival of 6-month-old spats. Animals were randomly collected from the hatchery, assigned to independent experimental tanks (n = 30 tank^-1^, mean individual Fresh Weight [_ind.FW_]= 0.13 ± 0.06 g), and allowed to acclimate for four days. Then, animals were exposed to a gradual warming [range 0.7-4.3 °C.h^-1^], reaching their respective randomly-assigned temperature treatments within 6 hours (to mimic the low-tide duration period). Oysters were then maintained at the target temperatures until mortality in the tanks reached 50 % (i.e., LT_50_). Oyster status (alive/dead) was checked every 30 minutes during the initial 24 h, and every 2-3 h thereafter. Assessment was done visually and by mechanical stimulation (gently poking the mantle). When animals did not respond to stimuli (valves were opened and did not respond to contact), time was recorded and the individual was removed from the tank. The experiment was stopped after 30 days, and duplicated over time.

#### 2.1.3 Thermal performance curve of juveniles (E3)

We examined the effect of 10 temperatures, ranging from 27 to 36 °C (1 °C increments; duplicated), on the survival sub-lethal physiological performance of 6-month-old animals. Spats were assigned randomly to independent experimental tanks (n = 250 tank^-1^, mean _ind.FW_ = 0.70 ± 0.50 g), and allowed to acclimate for 30 days. Once the acclimatory phase completed, animals were exposed to a gradual cooling [0.2 °C.h^-1^] /warming [range 0.2-1.3 °C.h^-1^] of the seawater within 6 hours. After 7 days of exposure, we estimated the scope-for-growth (*SFG*) of 6 individuals per treatment, by measuring respiration and ingestion rates using a closed-system respirometry approach (e.g., see Fly & Hilbish, 2013; detailed equations and energetic values conversion in **Supplementary information**).

### 2.2 Statistical analyses

All data analyses and modelling were done using R version 4.3.1 (R Core Team, 2023). Thresholds of statistical significance (α) were set at 0.05, unless specifically stated.

#### 2.2.1 Survival curves

In the experiment E1, exposure durations were not sufficient to identify directly LT_50_ (i.e., the exposure duration at which probability of mortality is equal to 50%), and implement the TDT curve. Therefore, as a first step, relative larval survival (E1; *warm* conditions) was plotted against time of exposure and fitted to a temperature-specific logistic regression model (generalized linear model, binomial distribution). The LT_50_ was then estimated using the car R package.

#### 2.2.2 Thermal performance curves (TPCs)

Mean survival data (E1) were modelled in relation to temperature (glm, binomial distribution) for the (i) embryogenesis and (ii) pelagic larval-development phases, separetly. In turn, spat metabolic rates (E3) were modelled as a thermal performance curve using univariate regressions (lm, normal distribution). We computed various models including linear, quadratic and cubic effects of temperature, and then compared their skill based on likelihood-ratio tests (LRT) and model outputs comparison (Akaike and Bayesian information criteria). Finally, residuals of the selected models were checked, using the DHARMa R package and diagnostic plots (Residuals *vs*. Fitted, QQ-plot, Scale-Location, Residuals *vs*. Leverage), for binomial and normal distributed models, respectively.

#### 2.2.3 Thermal death time curves (TDT)

For each life stage, measured elapsed times to reach 50 % mortality (i.e., *LT*_*50*_; in minutes, log_10_-transformed) were plotted against the exposure temperature (*temp*; in °C). TDT curves were then generated by fitting ordinary linear regressions (Rezende et al., 2014):

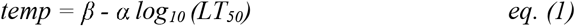

with β the intercept and α the slope. The TDT metrics (Rezende et al., 2014), namely the upper thermal limit for a specific duration of exposure (*t*; in min), *CTmax(t)* (°C), and the sensitivity (z) are calculated from the linear model output parameters:

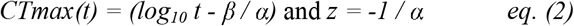

Because exposure durations differed between experiments E1 and E2, and not sufficient to identify the TDT breakpoint temperature (Jørgensen et al., 2021), we estimated Tc as the maximum temperature which did not induce acute heat failure.

#### 2.2.4 Estimation of cumulative thermal injury

The TDT parameters were then used to estimate the life stage-specific cumulative thermal injury expected under natural, fluctuating temperature conditions. Accumulation of thermal stress (condition Tenv > Tc) were calculated as a function of time, using free access R-scripts (https://github.com/MOersted/Termal-tolerances), based on the equation described in Jørgensen et al., (2021):

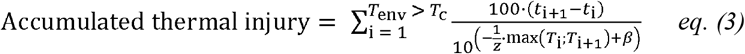

We investigated cumulative thermal injury at three contrasting sites across French Polynesia based on *in situ* temperature records. Nuku Hiva (140° 05’ W, 8° 51’ S) represents the most extreme site, where *P. margaritifera* inhabits tide pools with temperatures varying between 26 to >34 °C within a day (Reisser et al., 2019). Then, the atoll of Reao (136°37’ W, 18°51’ S) is a relatively warm lagoon due to its shallow depth and the limited exchange of water with the open ocean. Finally, the closed atoll of Takapoto (145°21’ W, 14°63’ S) represents a control site, notably due to its low seasonal variations (∼ 5 °C, recorded in 2021) and active pearl farming acitivty. To estimate the cumulative thermal injury at each of these sites, we used seawater temperature recorded during the warmest months. Measurements were done over eight days using iButton thermal loggers at Nuku Hiva (02 - 10/03/2023) and Reao (02 - 10/03/2023), and extracted from Liao *et al*. (2023) for Takapoto.

## 3. Results

### 3.1 Modelling and estimating thermal thresholds

#### 3.1.1 Survival curves

For the *warm* conditions of 36 and 38 °C, all larvae died within 180 minutes. These treatments were thus not considered in the subsequent analyses. All (binomial fixed effect) models for *warm* conditions showed a negative effect of exposure duration on larval survival. The 50% mortality (i.e., LT_50_) at 30, 32, and 34 °C was estimated at 1614, 1037 and 155 minutes, respectively.

#### 3.1.2 TDT curves

Linear regressions of log_10_(LT_50_) against temperature were generated for each life stage. Curves had a high coefficient of determination (R^2^) ranging from 0.89 to 0.94 (**Table 1**). TDT curves revealed that juveniles exhibited a higher CTmax(1h) (42.3 °C) than larva (36.1 °C). Thermal sensitivity (*z*) was also higher for the early developmental stage 3.36 °C than for juvenile spats (2.75 °C). The effects of temperature and life stage on LT_50_ were examined in a two-way ANOVA. Analysis results confirmed the significant effect of temperature and life stage on LT_50_ (*p* > 0.001), but no evidence of an interaction was detected (*p* = 0.23).

**Table 1.** **Parameters of the TDT curves** for the early-life (1 - 24 hpf) and later-life stage (6 months-old spat) of *Pinctada margaritifera*. Total thermal injury (%) accumulated over a 8 days period, based on natural small-scale thermal varitions at three sites in French Polynesia.

| Life-stage | TDT parameters |  |  |  | Cumulative thermal injury (%) |  |  |
| --- | --- | --- | --- | --- | --- | --- | --- |
|  | CTmax(1h) | Thermal sensitivity (z) | R<sup>2</sup> | est. Tc | Nuku Hiva | Reao | Takapoto |
| Early | 36.1 | 3.36 | 0.89 | 29 | 100 | 100 | 5.1 |
| Later | 42.1 | 2.75 | 0.94 | 34 | 30.2 | 1.2 | 0 |

#### 3.1.3 Integrative thermal tolerance landscape

The integrative framework, combining life stage-specific physiological and survival data, is presented in **Figure 1**. All thermal performance curves were best characterised with a quadratic function, except for respiration rate in spats (best fit :linear effect). The temperature of optimal performance (Topt) was 26.3 and 28.1 °C for embryos and D-shaped larva, respectively. In turn, the spat Topt for ingestion rate, and scope for growth were 29.8 and 29.6 °C, respectively. By overlapping the physiological performance curves (permissive range) with TDT curves (stressful range), we estimated Tc at 29 and 34°C, for the early-(larva) and later-(spat) life stages, respectively.

**Figure 1.**
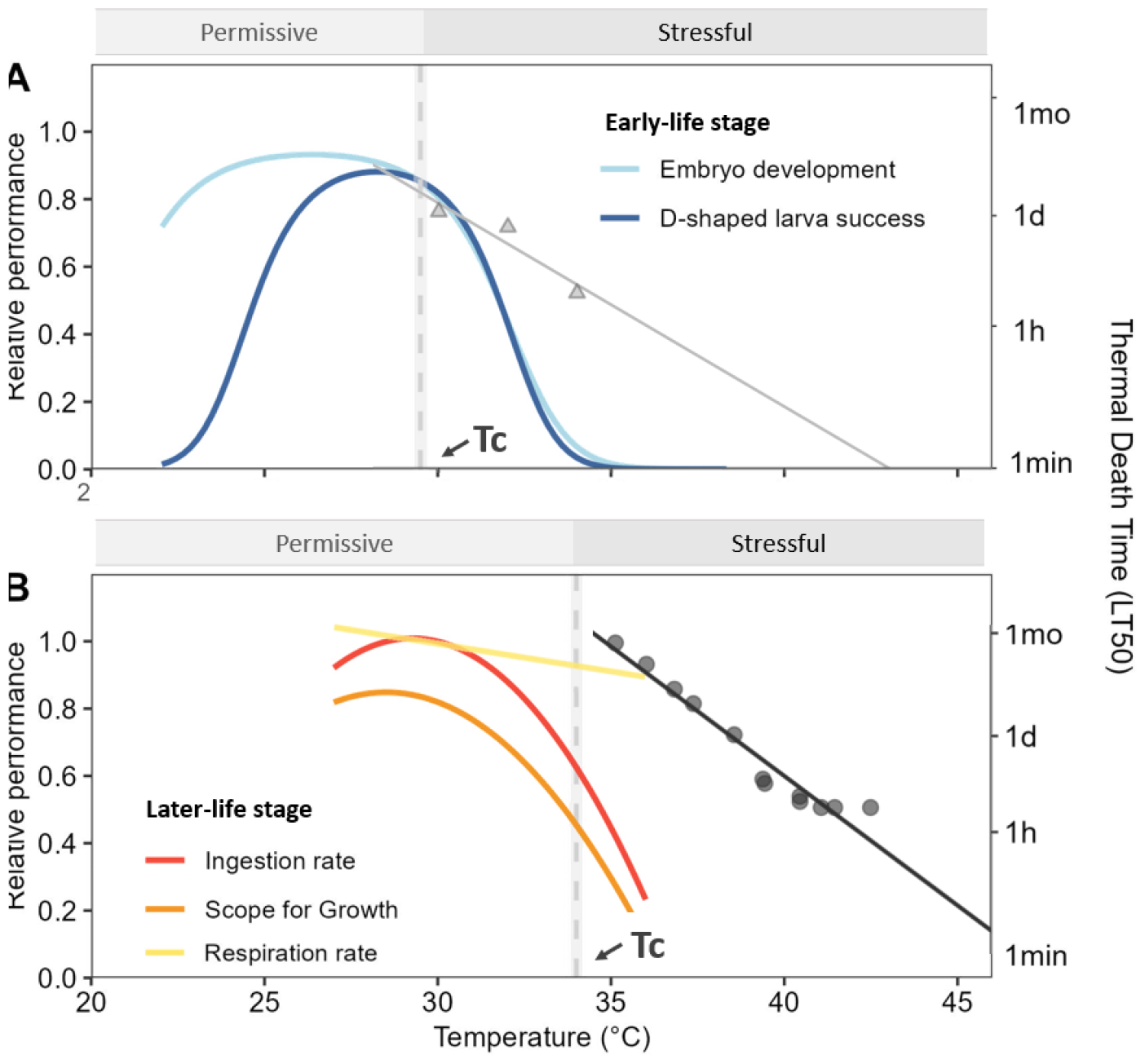
***Pinctada margaritifera* Thermal Tolerance Landscape for (A) Early** (1-24 hpf) and **(B) Later** (6 months-old spat) life stages. Thermal performance curves are fitted to relative mean performance (left y-axis) through binomial and linear regression models, respectively. TDT curves (right y-axis) are fitted to log_10_-transformed time causing a 50 % mortality (LT_50_) for pelagic larval development phase (grey triangles), and spats (black dots). TDT parameters can be found in Table 1. The critical temperature (Tc; grey dashed line), is shown to indicate the *stage-specific* transitional zone between the permissive and stressful temperature range.

### 3.2 Injury accumulation in natural fluctuating environments

We then used the thermal tolerance metrics estimated here to calculate the thermal injury accumulated by larvae and spats under ecologically realistic, *in situ* conditions. Sea-water temperature measurements (8 days), confirmed contrasting sites over the French Polynesian territory (**Figure 2**). Nuku Hiva showed the most extreme thermal profiles (mean 29.13°C□±□2.50LSD, min.: 24.10°C, max.: 42.40°C), while Reao mean 30.50°C□± □0.31□SD, min.: 29.74°C, max.: 31.52°C) and Takapoto (mean 28.46°C□± □1.05□SD, min.: 25.65°C, max.: 31.39°C) revealed to be the lagoon with the highest and the lowest mean temperature, respectively. Based on these divergent thermal regimes, results of cumulative injury varied with ontogeny and site (**Table 1**). Larvae accumulated more thermal injury than spats at every site. Based on natural thermal regimes recorded at Nuku Hiva (tide pools) and Reao (shallow lagoon), larvae reached 100% of accumulated injury (i.e., heat failure/death of the organism) within 840 and 1350 minutes, respectively. At the pearl-farming lagoon of Takapoto, larvae accumulated 5.1% injury within 8 days. Later-life spats only exhibited accumulated injury when exposed for 8 days to the thermal profiles of Nuku Hiva (30.2%) and Reao (1.2%); while no thermal injury was detected based on Takapoto’s *in situ* sea-water temperature.

**Figure 2.**
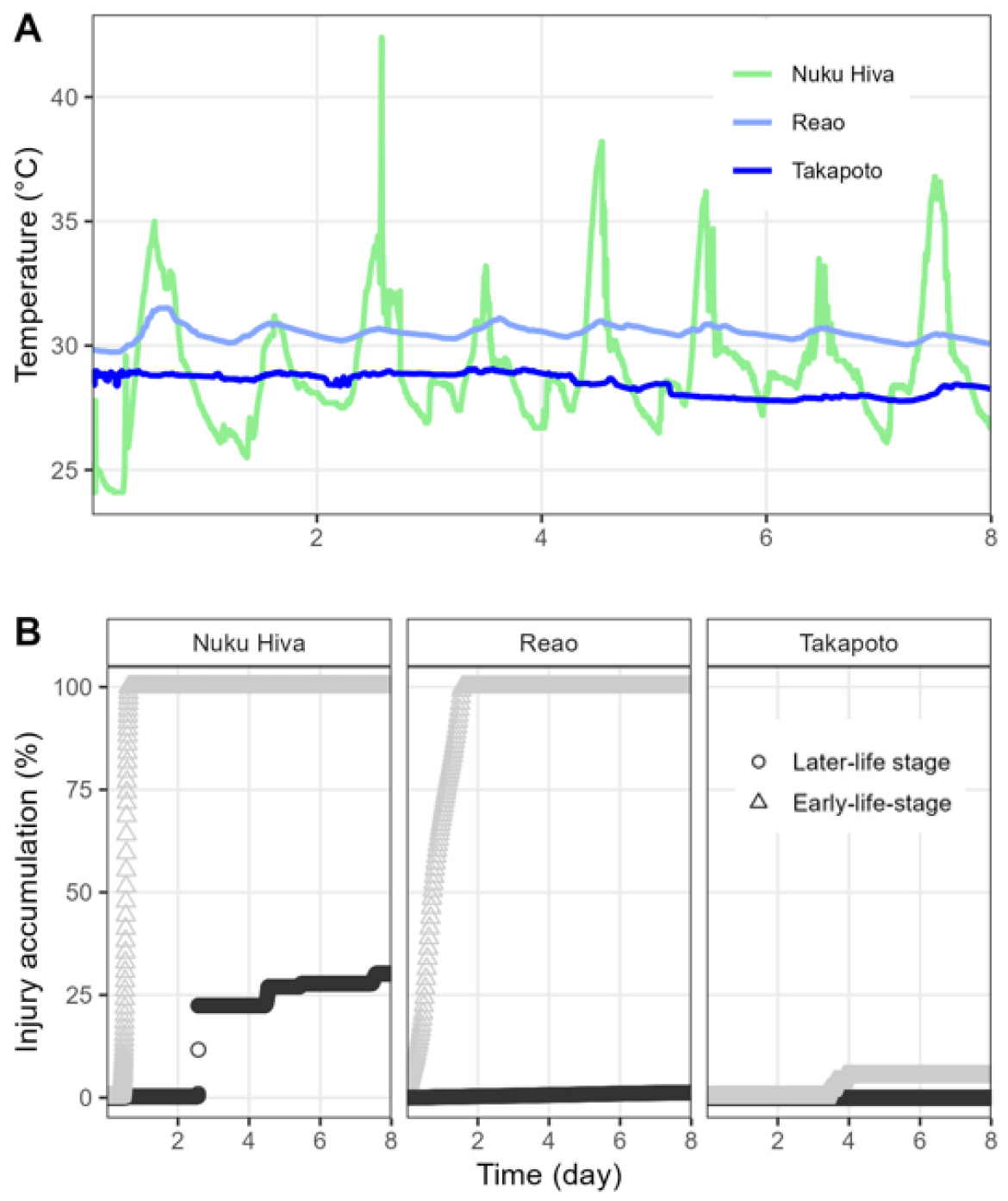
**(A) Natural temperature fluctuations** recorded *in situ* during the warmest months (March) at three sites, showing different thermal regimes. **(B) Stage-specific predicted accumulated injury** (see the formula in section 2.2.4.) for the two life stages: spats (black circles) and larva (grey triangles), throughout a week (8 days). 100% injury accumulation equals to heat failure (i.e., death of the organism).

## 4. Discussion

### Critical thermal limits

Using *Pinctada margaritifera* as a model species, we have quantified heat failure and performance under a broad range of temperatures, which allowed a fine-scale characterization of the species tolerance landscape, and a clear-cut estimation of the critical temperature (Tc) delimiting the permissive from the stressful temperature range.

By working with two life stages (early planktonic *vs*. spat), our study revealed an ontogenetic shift in lethal (CTmax) and sub-lethal (Tc) thermal limits, with higher vulnerability for early (planktonic) life stages, than for 6-month juveniles. Our results are in line with numerous studies on marine ectotherms; e.g., in fish (Dahlke et al., 2020), shallow-water echinoids (Collin et al., 2021), or tubeworms (Rebolledo et al., 2020). Different hypotheses have been proposed to explain the observed life stage-dependent thermal tolerance; e.g., oxygen-limitation due to the ventilatory and cardiorespiratory system’s development (Pörtner, 2002), or allometric constraints (Pörtner & Farrell, 2008). By providing such quantitative means, it now seems possible to further explore thermal tolerance at numerous life stages (from fertilization to spawners), and confirm such hypotheses. Ultimately, our results suggest that accurate predictions of population dynamics and evolutionary bottlenecks under ongoing climate change must consider nuances of species fundamental thermal niches, including the thermal limits of these early develpmental stages.

### In situ thermal injury accumulation

Translating laboratory observations of organisms’ heat stress responses to the natural environment remains challenging, in part, because the duration and intensity of thermal stress events are often unpredictable in the wild. The recent experimental validation that thermal injury is an additive process (above Tc) has provided an unprecedented quantitative tool for explicitly accounting for thermal heterogeneity in nature, thus improving our ability to assess species vulnerability to climate change (Jørgensen et al., 2021b).

We used this approach to evaluate the severity of thermal stress exhibited by larva and juvenile *P. margaritifera* experiencing three different natural fluctuating thermal regimes. As expected, bivalves accumulated high levels of thermal injury (100% for larvae, and 30% for spats) when exposed to the most ‘extreme’ site (Nuku Hiva; T°C > 38°C). Admittedly, this framework does not account for recovery ability (when T_env_ fluctuate between permissive and stressful range temperature) which might overestimate injury accumulation (Ørsted et al., 2022b). The conceptual underpinnings of this temperature and duration-dependent repair function remain elusive, in part due to inherent difficulties associated with rapid hardening (i.e., transient response conferring enhanced heat tolerance following a sub-lethal exposure) physiological processes (Ørsted et al., 2022b). Still, considering the short time of exposure calculated to induce heat failure in larvae (12 and 22 h for Nuku Hiva and Reao, respectively), we expect that recovery period might be limited and insufficient to prevent high selective pressure on early stage. In addition to the high degree of mortality resulting from the transition from a pelagic to benthic stage (Jenkins et al., 2009), we expect a important reduction of settlement during the warmest months at these sites. The significant match between the estimated critical temperature (Tc ∼29°C) and the mean annual sea-surface temperature in Takapoto Atoll (28.3 ± 0.8 °C) suggest that species are living closer to their upper thermal limits than previously estimated (Le Moullac et al., 2016). Overall, this approach provides (i) the foundations for comparing the species fundamental *vs*. realized thermal niches, which can help investigating evolutionary and biogeographic ecological processes, and (ii) a promising tool to quantify the impacts of extreme events (e.g., marine heat waves).

## 5. Conclusion

The present study reinforces the importance of defining ontogeny-specific thermal tolerance limits, to avoid underestimating the vulnerability of individuals and populations threatened by global warming. Indeed, thermal stress assays done using *P. margaritifera* revealed an ontogenetic shift in lethal (CTmax) and sub-lethal (Tc) thermal limits, with higher vulnerability for early-life (planktonic) stages, than for 6-month spats.

By integrating these estimations of thermal limits and the thermal injury accrued additivly over time (Jørgensen et al., 2021a), we provide predictions of heat-failure risk for three contrasting sites in French Polynesia. Results obtained for relatively stable thermal regime’s atolls (Reao and Takapoto), were consistent with specie’s natural observations (absence and presence, respectively). However, injury calculated for the specific tidal habitat of Nuku Hiva (i.e., Tenv crossing permissive and stressful temperature ranges) inducates thermal stress in spats, and high selective pressure on early stage during warmest months.

Still, the exploration of the full tolerance landscape of organisms offers possibilities for more advanced mechanistic explorations. Indeed, sublethal effects at high temperature (e.g., oxidative stress, depletion of energy reserves, infertility, etc) are expected to increase thermal sensitivity with longer exposure (Kingsolver & Woods, 2016), resulting lower growth and reproductive output. Indeed, predicted elevation of temperature will not necessarily induce mortality (presence or absence), but may rather induce nuanced changes in population abundance, geographic range limits and/or work as a driver of local adaptation. Such impacts may be particularly important for climate-sensitive economic sector – such as molluscan shellfish farming (Fly & Hilbish, 2013), and specially those dependent on wild-spat collection (Doubleday et al., 2013). Combining tools capable of quantifying such sublethal effects (e.g., Thomas & Bacher, 2018), recovery effectiveness (Jørgensen et al., 2021b), as well as species thermal limits - would be a major step forward for prediction accuracy, and the guidance of resource management and conservation programs.

## Aknowledgement

We thank Lauriane Bish for support in animal rearing and algae production. Reao temperature data collected within the GAIA project funded by the Agence National de la Recherche [ANR-21-CE32-0011-01 GAIA] were kindly provided by S. Van Wynsberge and V. Teaniniuraitemoana. This work is part of the PhD thesis of K.L and supported by the PinctAdapt project.

## Additional files

**Additional file 01**: Supplementary information

