## Supplementary Information for "Exploring thermal tolerance across time and space in a tropical bivalve, *Pinctada margaritifera*"

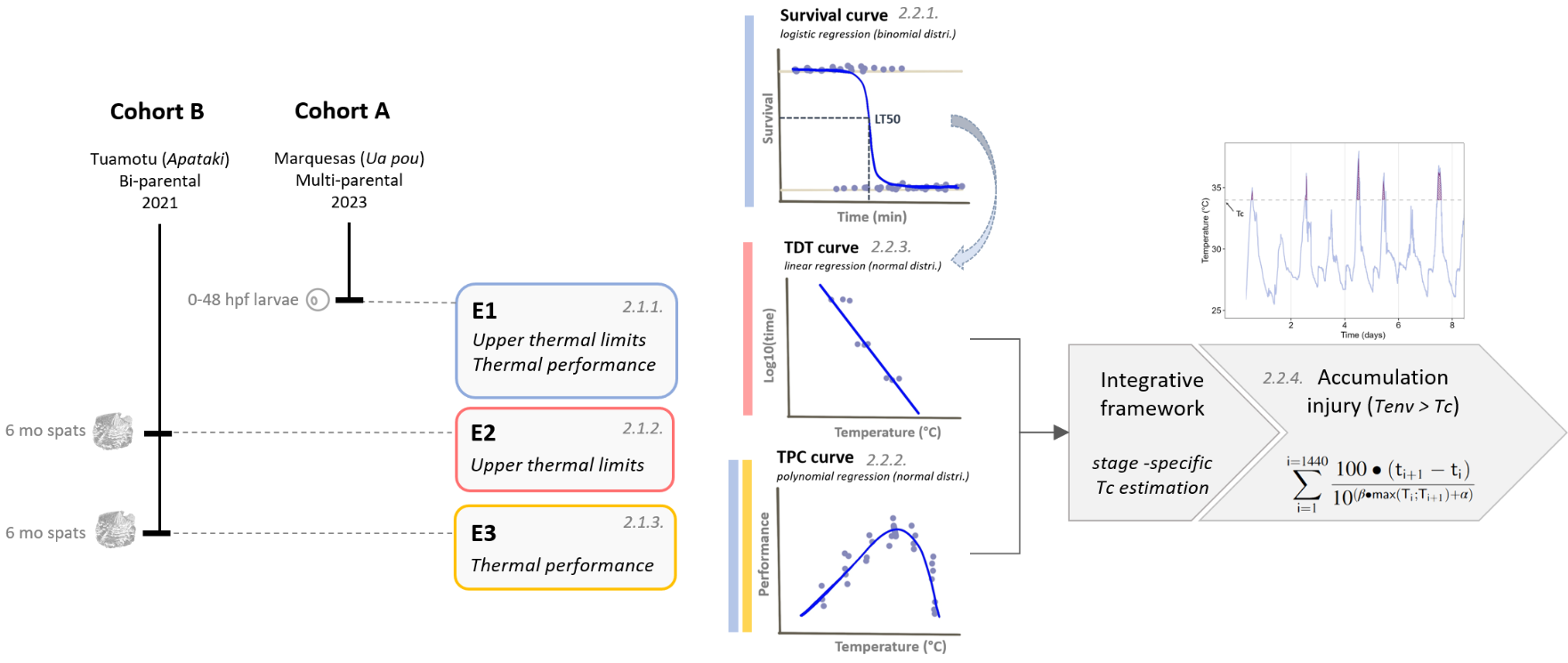

**Figure S1. Graphical abstract.** Three different experiments (E1, E2 and E3) were performed using hatchery-produced individuals issued from two reproduction events (see below cohorts A and B in **Note S1**). The E1 experiment aimed to assess larval development completion (i.e., performance and upper limits). In turn, the E2 and E3 allowed investigating the thermal sensitivity of spats experiencing stressful and permissive temperatures, respectively.

**Note S1. Animals.** Cohort A. Wild parents (4 males x 2 females) were collected in January 2023 at the Marquesas archipelago (Ua Pou island, 9°22′60″S, 140°03′W) and transferred to Ifremer’s marine concession in Vairao (Tahiti, French Polynesia). On April 2023 May 2021, breeders were transferred from the natural lagoon to the controlled experimental facilities for six weeks, in order to initiate a synchronized gametogenesis. Breeders maintenance, spawning induction and larval rearing were performed as described in Hui et al. (2011) and Ky et al. (2013). Released gametes from each breeder were immediately isolated in beakers, and quality controlled (visual egg appearance and sperm mobility) before pooling.

Cohort B*.* Wild parents (1 male x 1 female) were collected on October 2015 at the Tuamotu archipelago (Apataki atoll, 17°48′31.7″S, 149°17′41.4″W). In April 2023, breeders entered a six-week conditioning period. Conditioning, spawning triggering and fertilization were done as for cohort A. Progenies were further maintained in common-garden, under hatchery-controlled conditions, until the experiments began (i.e., 6 months later; E2 and E3). This F1 full-sib family strategy was selected (i) to avoid confounding effects as a result of thermal environment history experienced by the wild broodstock (e.g., carry-over effects), and (ii) in a way to reduce the genetic background, ontogenetic and physiological condition variations within individuals (i.e., making more comparable the observed phenotypes).

**Note S2. Experimental tanks*.*** The experimental -controlled set up consisted of 29.2-L independent open flow tanks, supplied by 10-µm filtered and UV-treated seawater (flow rate = 20-L .h^-1^, turnover = 68% .h^-1^). Food was regulated by an external microalgae input (1:1 mix of *Tisochrysis galbana* and *Chaetoceros gracillis*), maintained at 800 μm3 mL^−1^ cells at the outlet of the tank, and daily controlled using an electronic particle counter (Multisizer3, Beckman Coulter; 100-μm aperture tube). Oxygen stones allowed a well-oxygenated environment, and homogeneous concentration of phytoplankton cells. Overall, maintenance conditions were recreating ambient sea water parameters (temperature 27°C, salinity 36 psu, pH_NBS_ 8.3, DO_2_ saturation 95%, photoperiod 14:10 D:L), which were daily checked (multi-Parameter WTW®). For thermal stress assays, seawater temperature was regulated by means of heat resistance (IKS^®^ Aquastar system) or chiller (TECO^®^), for heat or cold conditions, respectively.

**Note S3. Metabolic measurements and calculations**. Spats were randomly sampled from experimental tanks, and placed in individual closed-system respirometry chambers (120 mL volume), fulfilled with filtered and oxygen-saturated seawater, and maintained at temperature relating to the treatment assigned. After the bivalves appeared to have started to filter (as indicated by open valves and the mantle edge and tentacles in an outstretched position), respiration measurements started, for a total duration of 1.5 hours. Measurements were done by means of an optical oxygen sensor coupled with an Opto-F1 UniAmp (Unisense®). Once oxygen measurements ended, around 1.8.10^5^ cells of phytoplankton^[1]^ were added in the sealed volume, and waited for another 1.5 hours before sample 10 mL seawater. The final phytoplankton concentration was measured using an electronic particle counter (Multisizer3, Beckman Coulter; 100-μm aperture tube). Ingestion Rates (**IR**; cell .h^-1^) and Respiration Rates (**RR**; mg O_2_ .h^-1^) were converted to a standard animal basis, using the formula:

*Ys = (Ws/We)b × Ye	(eq. S1)*

Where Ys is the standard metabolic rate, Ws is the standard dry weight (1 g), We is the measured dry weight, Ye is the measured physiological activity, and b is the allometric coefficient. Based on Savina & Pouvreau (2004), we used b allometric coefficient of 0.66 for IR and 0.75 for RR. Then Scope for Growth (**SFG**) was calculated according to the following equation :

*SFG = (IR x AE) – RR	(eq. S2)*

where *AE* is the organic matter assimilation efficiency (%). The values of 20.3 J for 1 mg of particulate OM (Bayne et al., 1987) and 14.1 J for 1 mg O_2_ (Bayne & Newell, 1983; Gnaiger, 1983) were used for to convert metabolic rates into energetic values. For the AE (Widdows & Staff, 2006), the value of 0.45 was used, which does not vary with temperature in *P. margaritifera* (Le Moullac et al., 2016).

**Table S1**. Linear, quadratic and cubic effects of temperature on successful completion of early life stages : embryo (1 - 6 hpf) and larval (1 - 24 hpf) development. Estimates are from a binomial mixed-effects regression model. The model selection was performed by comparing fits of models, by means of Akaike Information Criterion (AIC) and likelihood-ratio tests (LRT; comparison value with the upper line model).

| **life stage** | **Fixed effects** | **AIC** |  | **LRT** | | | |
| --- | --- | --- | --- | --- | --- | --- | --- |
|  |  |  |  | **χ2** | **d.f.** |  | ***P*** |
| Embryo development (1 - 6 hpf) | Temperature (linear effect) | 167.95 |  |  |  |  |  |
|  | Temperature (quadratic effect) | 140.61 |  | 29.35 | 1 |  | < 0.001 |
|  | Temperature (cubic effect) | 139.83 |  | 2.77 | 1 |  | 0.095 |
| Larval developement (1 - 24 hpf) | Temperature (linear effect) | 169.45 |  |  |  |  |  |
|  | Temperature (quadratic effect) | 127.50 |  | 43.95 | 1 |  | < 0.001 |
|  | Temperature (cubic effect) | 129.44 |  | 0.06 | 1 |  | 0.808 |

**Table S2.** Linear, quadratic and cubic effects of temperature on between measured spat’s metabolic parameters : Ingestion, Respiration and Scope for Growth. Estimates are from a linear fixed-effects regression model. Summary of the Akaike and Bayesian Information Criterion (AIC. BIC), residual sum of squares (RSS) and R-squared (R^2^). The regression model revealing the minimum AIC and BIC value was considered as the best fit.

| **Metabolic rate** | **Model** | **AIC** | **BIC** | **RSS** | | **R^2^** |
| --- | --- | --- | --- | --- | --- | --- |
| Ingestion | Temperature (linear effect) | 7,28 | 8,19 | 0.665 | | 0.069 |
|  | segmented | 2,19 | 3,70 | 0.268 | | 0.369 |
|  | Temperature (quadratic effect) | 2,11 | 3,32 | 0.324 | | 0.223 |
|  | Temperature (cubic effect) | 2,42 | 3,94 | 0.274 | | 0.284 |
| Respiration | Temperature (linear effect) | -1,76 | -0,85 | 0.269 | | 0.069 |
|  | segmented | -1,64 | -0,13 | 0.182 | | 0.369 |
|  | Temperature (quadratic effect) | -1,56 | -0,35 | 0.225 | | 0.223 |
|  | Temperature (cubic effect) | -0,38 | 1,13 | 0.207 | | 0.284 |
| Scope For Growth | Temperature (linear effect) | 12,42 | 13,33 | 1.112 | | 0.405 |
|  | segmented | 11,70 | 13,22 | 0.694 | | 0.629 |
|  | Temperature (quadratic effect) | 10,95 | 12,16 | 0.786 | | 0.579 |
|  | Temperature (cubic effect) | 12,62 | 14,13 | 0.761 | | 0.593 |

1. Based on pilot experiments
